# Kestrels of the same colony do not overwinter together

**DOI:** 10.1101/2022.12.14.520401

**Authors:** Jorge García-Macía, Munir Chaouni, Sara Morollón, Javier Bustamante, Lina López-Ricaurte, Juan Martínez-Dalmau, Beatriz Rodríguez-Moreno, Vicente Urios

**Affiliations:** Grupo de Investigación Zoología de Vertebrados, Universidad de Alicante, Apdo. 99, E-03080 Alicante, Spain; Department of Wetland Ecology, Estación Biológica de Doñana (CSIC), C/ Américo Vespucio 26, 41092, Sevilla, Spain; GREFA, C/ Monte del Pilar S/N, E-28220 Majadahonda, Madrid, Spain

**Keywords:** *Falco naumanni*, raptor, spatial ecology, movement ecology, GPS telemetry, wintering, non-breeding

## Abstract

Coloniality is one of the most common strategies in birds. While the lesser kestrel (*Falco naumanni*) is a colonial raptor during the breeding period, it is not known whether individuals from the same breeding colony aggregate during the non-breeding period too. We GPS-tracked 40 adult lesser kestrels from different Spanish breeding colonies to study the degree of spatial aggregation between individuals from the same breeding colony in their West African non-breeding range. Lesser kestrels in our study used a large area from a wide longitudinal strip in the western Sahel: individuals used 143,697 ± 98,048 km^2^ on average during the entire non-breeding period (95% KDE), and 1,359 ± 1,424 km^2^ per week. On the other hand, the individuals traveled 6205 ± 2407 km on average during the entire non-breeding period, and 41.1 ± 11.8 km per day. There were no differences between the sexes in any of those variables. Individuals from the same breeding colony were not aggregated during the non-breeding period because the overlap between their areas (38.8 ± 21.4 %) was not higher than that randomly expected. In conclusion, our study reveals some aspects of the non-breeding spatial ecology of the lesser kestrel, allowing a better understanding of the relationship of colonial birds out of the breeding season.

## 1. Introduction

Coloniality is one of the most common strategies in birds. Up to 13% of birds’ species cluster in colonies during the breeding period (Lack 1968). Colonial birds share common grounds and tasks during some stages of their life-cycle, with a balance between costs and benefits (Rolland *et al*. 1998). Costs involve competition for food or nesting sites, and an increased risk of transmission of diseases and cannibalism (Wittenberger and Hunt 1985, Birkhead and Møller 1992). Benefits involve information sharing among individuals regarding foraging areas, nesting selection, or anti-predator defense (Ward and Zahavi 1973, Clode *et al*. 2000, Arroyo *et al*. 2001, Barta and Giraldeau 2001, Danchin *et al* 2004, Aparicio *et al*. 2007). Colonial breeding can sometimes offset the benefits of a territorial behaviour, due to the advantage of information on food location (Rolland *et al*. 1998). Coloniality may be specially useful during high-demanding energy periods like breeding (Di Maggio *et al*. 2013), when pairs must dedicate most of the resources to feed and protect the chicks.

The different species of raptors can adopt different spatial strategies throughout their annual cycle. Some long-lived raptors follow a territorial strategy through the entire year (Pérez-García *et al*. 2013, Sur *et al*. 2020, Grainger Hunt *et al*. 2021). On the other hand, many species form aggregations of tens or thousands of individuals during the breeding period (Arroyo *et al*. 2001, Cecere *et al*. 2018) or the non-breeding period (Senar and Borras 2004, Arroyo and García 2007, Pilard *et al*., 2011, Urios and García-Macía 2022). However, little is known about the connectivity of the breeding colonies during the non-breeding (or wintering) period. Do the breeding neighbors also share non-breeding grounds far away from their nests? The marked changes regarding physiological requirements, environmental characteristics and threats between the breeding and non-breeding periods entail that individuals change their foraging and spatial strategy to be efficient and survive (Urios *et al*. 2017, López-López *et al*. 2021, Urios and García-Macía 2022), so it is necessary to study how individuals are spatially related during the entire year.

The lesser kestrel, an insectivorous raptor which follows a colonial strategy during the breeding period, may be an interesting model species to study the shift in the degree of aggregation between the breeding and non-breeding periods. Regarding the breeding season, some studies reveal a great degree of overlap between the home ranges of lesser kestrels from the same colony, even with strict spatial segregation of those home ranges between neighboring colonies (Cecere *et al*. 2018). However, information about how individuals from the same breeding colony overlap during the non-breeding period in West Africa is still being resolved. As noted above, it is frequent among birds, including the lesser kestrel, the use of communal roosts during winter, but the provenance and aggregation of the individuals arriving at these places are not always clear: they may comprise mainly individuals from the same breeding colony or be the result of mixed aggregations. Some previous works have studied the migratory connectivity of different lesser kestrel populations using a large-scale approach (Sarà *et al*. 2019), but there are no studies at a smaller scale on the spatial relationship of the individuals from the same breeding colony during the non-breeding period.

We analyze the data of 40 adult lesser kestrels from different Spanish breeding colonies to study the degree of spatial overlap among individuals from the same breeding colony during the non-breeding period in West Africa. Our objectives were: (1) to estimate the size of the nonbreeding grounds; (2) to determine the degree of overlap between individuals from the same breeding colony during this period, and compare with the random overlap between individuals from different breeding colonies.

## 2. Materials and methods

### Study species

The lesser kestrel (*Falco naumanni* Fleischer, 1818; Falconidae) is a small migratory raptor with a large geographical range in Eurasia and Africa. Their breeding populations extend from Central Asia to the Mediterranean region (Bijleveld 1974, Cramp and Simmons 1980, Ortego 2016, GBIF 2022). Individuals from the Eastern Palearctic usually follow a longer eastern migratory route, in order to spend the winter in East and South Africa (Ferguson-Lees and Christie 2001), while birds from the Western Palearctic follow a shorter western migratory route across the Mediterranean sea (Sarà *et al*. 2019), to reach their non-breeding grounds in sub-Saharan Africa. The diet of the lesser kestrel is essentially insectivorous, sometimes complemented with small mammals and reptiles (Rodríguez *et al*. 2006, Martín *et al*. 2007, Rodríguez *et al*. 2010). During the breeding period, this species prefers open areas like cereal pseudo-steppes (Atienza and Tella 2004), but it is also linked to urban environments and human constructions, where they often nest (Ortego 2016). It is a gregarious bird that usually forms colonies of dozens of pairs during the breeding season (Bustamante *et al*. 2021).

Globally, the lesser kestrel is considered as “Least Concern” because their entire population is currently “stable”, with 80,000-134,000 mature individuals (IUCN 2021). However, it has been considered “vulnerable” in Spain since the 1990s (Bustamante *et al*. 2021). In this country, around 10,000 breeding pairs have been estimated, distributed throughout more than 2,000 colonies and isolated breeding areas (Bustamante *et al*. 2020). The main threats for this species in Spain are the reduction of insects and other invertebrates in agricultural areas, restoration of buildings affecting nesting colonies, the wind and photovoltaic energy development in foraging areas, and different affections on non-breeding areas and migratory passage sites (Bustamante *et al*. 2021).

### Tagging and sample size

Forty adult lesser kestrels (21 females, and 19 males) tagged at different colonies in Spain (Table 1) provided data on non-breeding movements after returning to their breeding colony. We obtained data for 55 non-breeding seasons between 2016 and 2021.

**Table 1.** Metadata of the 40 lesser kestrels tagged with Solar GPS biologgers. Sex, age, colony (province), and beginning/end of tracking of each individual are shown.

| ID | Age | Sex | Colony (Spanish province) | Beginning of tracking | End of tracking |
| --- | --- | --- | --- | --- | --- |
| 16639 | Adult | Male | Baena (Córdoba) | 2017 | 2018 |
| 16643 | Adult | Female | Baena (Córdoba) | 2017 | 2019 |
| 16687 | Adult | Female | Castelló de Empuries (Gerona) | 2017 | 2018 |
| 16688 | Adult | Male | Castelló de Empuries (Gerona) | 2017 | 2019 |
| 17281 | Adult | Female | Tarancón (Cuenca) | 2018 | 2019 |
| 17292 | Adult | Female | Tarancón (Cuenca) | 2018 | 2019 |
| 17240 | Adult | Male | Villares del Saz (Cuenca) | 2018 | 2019 |
| 17245 | Adult | Female | Villares del Saz (Cuenca) | 2018 | 2019 |
| 17288 | Adult | Female | La Almarcha (Cuenca) | 2017 | 2019 |
| 16690 | Adult | Male | Can Viure (Barcelona) | 2017 | 2019 |
| 16611 | Adult | Male | Fraga (Huesca) | 2016 | 2019 |
| 17235 | Adult | Male | Arganda del Rey (Madrid) | 2018 | 2019 |
| 16127 | Adult | Male | Perales del Río (Madrid) | 2017 | 2019 |
| 16137 | Adult | Male | Perales del Río (Madrid) | 2017 | 2020 |
| 17256 | Adult | Female | Perales del Río (Madrid) | 2018 | 2019 |
| 17261 | Adult | Male | Perales del Río (Madrid) | 2018 | 2019 |
| 16275 | Adult | Female | Pinto (Madrid) | 2017 | 2018 |
| 16212 | Adult | Male | Pinto (Madrid) | 2016 | 2017 |
| 17239 | Adult | Female | Pinto (Madrid) | 2018 | 2019 |
| 17237 | Adult | Male | Pinto (Madrid) | 2018 | 2020 |
| 17241 | Adult | Female | Pinto (Madrid) | 2019 | 2020 |
| 17250 | Adult | Male | Torrejón de Velasco (Madrid) | 2018 | 2020 |
| 17251 | Adult | Male | Torrejón de Velasco (Madrid) | 2018 | 2019 |
| 17253 | Adult | Male | Torrejón de Velasco (Madrid) | 2018 | 2019 |
| 17254 | Adult | Male | Torrejón de Velasco (Madrid) | 2018 | 2020 |
| 17278 | Adult | Female | Navalcarnero (Madrid) | 2018 | 2020 |
| 17280 | Adult | Female | Navalcarnero (Madrid) | 2018 | 2019 |
| 17282 | Adult | Female | Navalcarnero (Madrid) | 2018 | 2019 |
| 17262 | Adult | Female | Navalcarnero (Madrid) | 2018 | 2019 |
| 16679 | Adult | Female | Villalpando (Zamora) | 2017 | 2019 |
| 16661 | Adult | Female | Villalpando (Zamora) | 2017 | 2018 |
| 16584 | Adult | Female | Villalpando (Zamora) | 2017 | 2018 |
| 16616 | Adult | Female | Villalpando (Zamora) | 2017 | 2017 |
| 17195 | Adult | Male | Espacio Natural de Doñana (Sevilla) | 2018 | 2019 |
| 17199 | Adult | Female | Espacio Natural de Doñana (Sevilla) | 2018 | 2019 |
| 17218 | Adult | Male | Espacio Natural de Doñana (Sevilla) | 2018 | 2019 |
| 17210 | Adult | Male | Silo de la Palma del Condado (Huelva) | 2018 | 2019 |
| 17214 | Adult | Female | Silo de la Palma del Condado (Huelva) | 2018 | 2019 |
| 17215 | Adult | Female | Silo de la Palma del Condado (Huelva) | 2018 | 2019 |
| 17219 | Adult | Male | Silo de la Palma del Condado (Huelva) | 2018 | 2019 |

Lesser kestrels were captured at the breeding colony, usually during courtship while roosting at the nest. After capture, individuals were ringed, weighed (mean = 146 g; n = 40) and measured. The sex was determined by morphological characteristics, given the sexual dimorphism of the species (Cade and Digsby 1982). A GPS-biologger transmitter was attached to the back of each individual by a back-pack harness tied with tefflon ribbon, designed to allow its release after a few years of monitoring. All the transmitters were Nano-GPS from PathTrack (Leeds, UK) model NanoFix GEO+RF (weigh = ~ 4g, less than 5% of birds’ weight), thus complying with the recommended standard (Kenward 2001).

Biologgers provided GPS fixes every 15 min to 1h from dawn to dusk during the non-breeding season, and downloaded data to a base located at the breeding colony. Some biologgers provided locations 24 hours a day from February to October, but night locations were excluded. Only kestrels that migrated back to the breeding colony provided data on non-breeding movements. Locations were filtered at homogeneous 1-hour frequency to avoid bias in subsequent calculations. Locations were transformed to UTM coordinates (WGS 84; EPSG: 32630).

### Traveled distances and area sizes

The beginning of non-breeding season was established when individuals arrived to the Sahel from Spain, that is, at the end of post-breeding migration. The end of non-breeding season was established when individuals left the Sahel, with the beginning of pre-breeding migration.

We calculated the hourly traveled distance as the Euclidian distance between two consecutive locations. All hourly distances calculated for each day were summed to obtain the daily traveled distance. We also calculated the mean daily traveled distance and the total traveled distance for the entire non-breeding season.

We estimated the size of the non-breeding areas using 95 and 50% Kernel Density Estimators (KDE). We used all locations of each individual and non-breeding season to estimate the global non-breeding area, and the locations of every week to estimate the weekly non-breeding area.

We used Linear Mixed Models (LMM) to test the differences in mean daily traveled distances and total traveled distances between sexes. “Sex” was established as a fixed effect, and “individual” and “non-breeding period” were set as random effects. We tested the normality of the residuals of the models using a Shapiro-Wilk test.

We used Mann-Whitney-Wilcoxon tests (MWW) to analyze the differences between sexes in the non-breeding area size (mean total KDEs and median weekly KDEs; data were non-normal).

### Overlapping between non-breeding areas

We analyzed the overlap between non-breeding areas (KDE 95%) of individuals from the same breeding colony during the same year (n=12) to determine whether individuals from the same breeding colony wintered in the same area. We also estimated the overlap between non-breeding areas of individuals from each colony and those from the rest of the colonies. A χ^2^ (chi-squared) test was performed to compare between those two variables. In addition, when data for at least two winters were available for the same individual (n=9), we calculated the repeatability of non-breeding areas as the year-to-year overlap of the non-breeding areas, following the same method as in the previous section.

All statistical analyses were performed with R Software v. 4.0.5. Maps were made with QGIS 3.16.6. A significant level was established at < 0.05.

## 3. Results

### Home range sizes and traveled distances

The 40 lesser kestrels spent the winter in the western Sahel (between 18° and 12° north latitude, and between −18° and −2° west longitude), occupying a longitudinal strip of about 1600 km (Figure 1).

**Figure 1.**
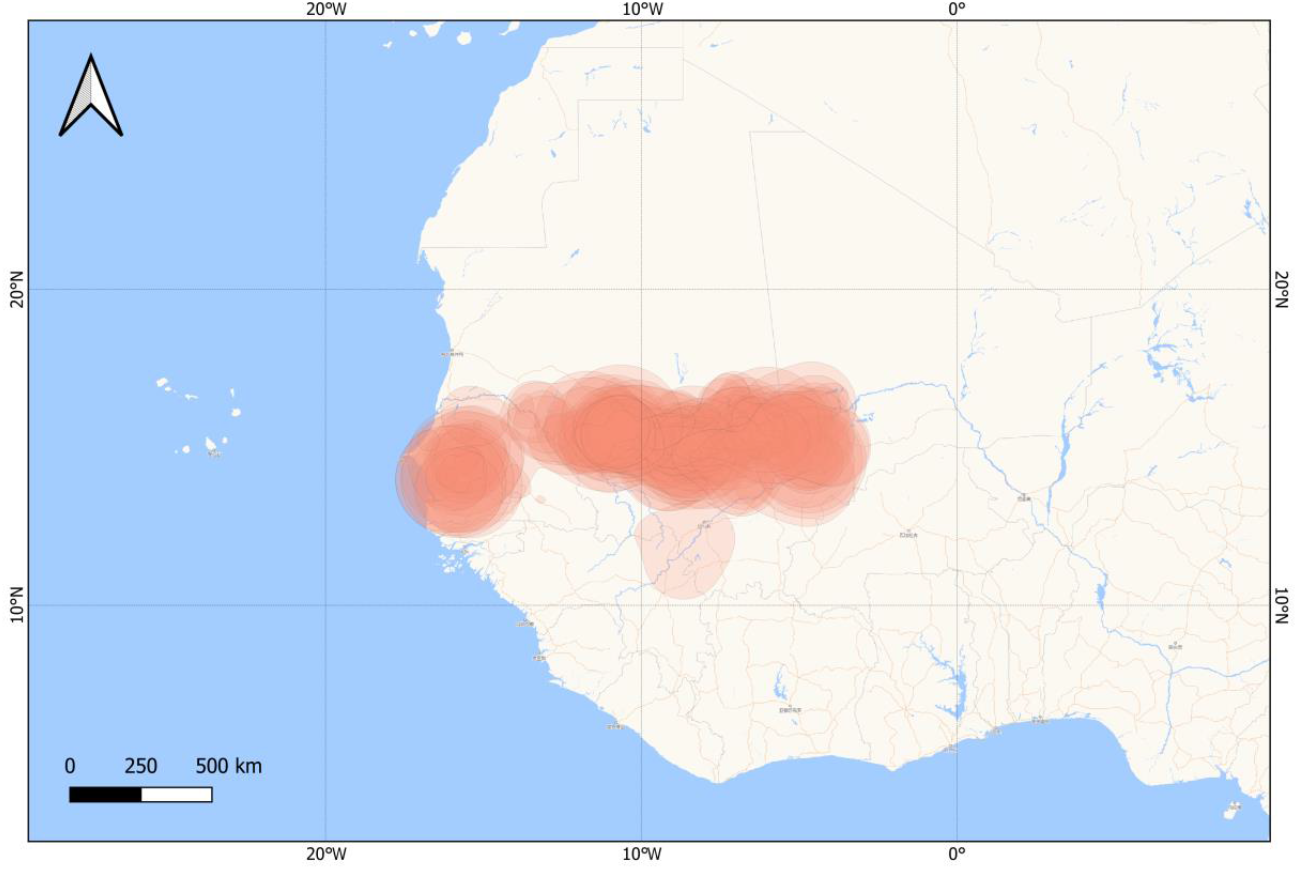
All non-breeding areas (KDE 95%) of the 40 lesser kestrels during 55 non-breeding periods.

There was a great variability in the size of the non-breeding areas (Table 2, Table S1). The lesser kestrels used, on average, 143,697 ± 98,048 km^2^ regarding 95% KDE, and 29,414 ± 20,572 km^2^ regarding 50% KDE (Table 2). The global area occupied was divided into smaller weekly areas between which individuals moved throughout their non-breeding period (Figure 2). The median for the weekly 95% KDEs was 1,359 km^2^’, and 248 km^2^ for the 50% KDEs (Table 2).

**Table 2.** Mean values of the non-breeding periods, regarding dates, duration, traveled distances and kernel density estimations (KDE). Values are shown as “mean ± *SD* (minimum-maximum)”, calculated by using mean values of each non-breeding season. Specific values are shown in Table S1.

|  | Non-breeding beginning | Non-breeding end | Days of non-breeding period | Total traveled distance (km) | Mean daily traveled distance (km) | Global 95% KDE (km<sup>2</sup>) | Global 50% KDE (km<sup>2</sup>) | Median weekly 95% KDE (km<sup>2</sup>) | Median weekly 50% KDE (km<sup>2</sup>) |
| --- | --- | --- | --- | --- | --- | --- | --- | --- | --- |
| Overall (n=55) | 29/08/18 | 24/01/19 | 147 ± 27 | 6205 ± 2407 | 41.1 ± 11.8<br>(20.1-64.5) | 143697 ± 98048<br>(5001-445393) | 29414 ± 20572<br>(750-96323) | 1359 ± 1424<br>(64-7359) | 248 ± 280<br>(7-1403) |
| Male (n=30) | 09/09/18 | 02/02/19 | 146 ± 25 | 5769 ± 2195 | 38.6 ± 10.4<br>(21.4-64.5) | 145253 ± 87939<br>(5001-279949) | 30020 ± 18709<br>(750-58163) | 1643 ± 1763<br>(76-7359) | 313 ± 352<br>(15-1403) |
| Female (n=25) | 16/08/18 | 12/01/19 | 149 ± 30 | 6727 ± 2587 | 44.0 ± 13.0<br>(20.1-63.3) | 141829 ± 110811<br>(12770-445393) | 28687 ± 22983<br>(2764-96323) | 1018 ± 767<br>(64-3105) | 169 ± 124<br>(7-464) |

**Figure 2.**
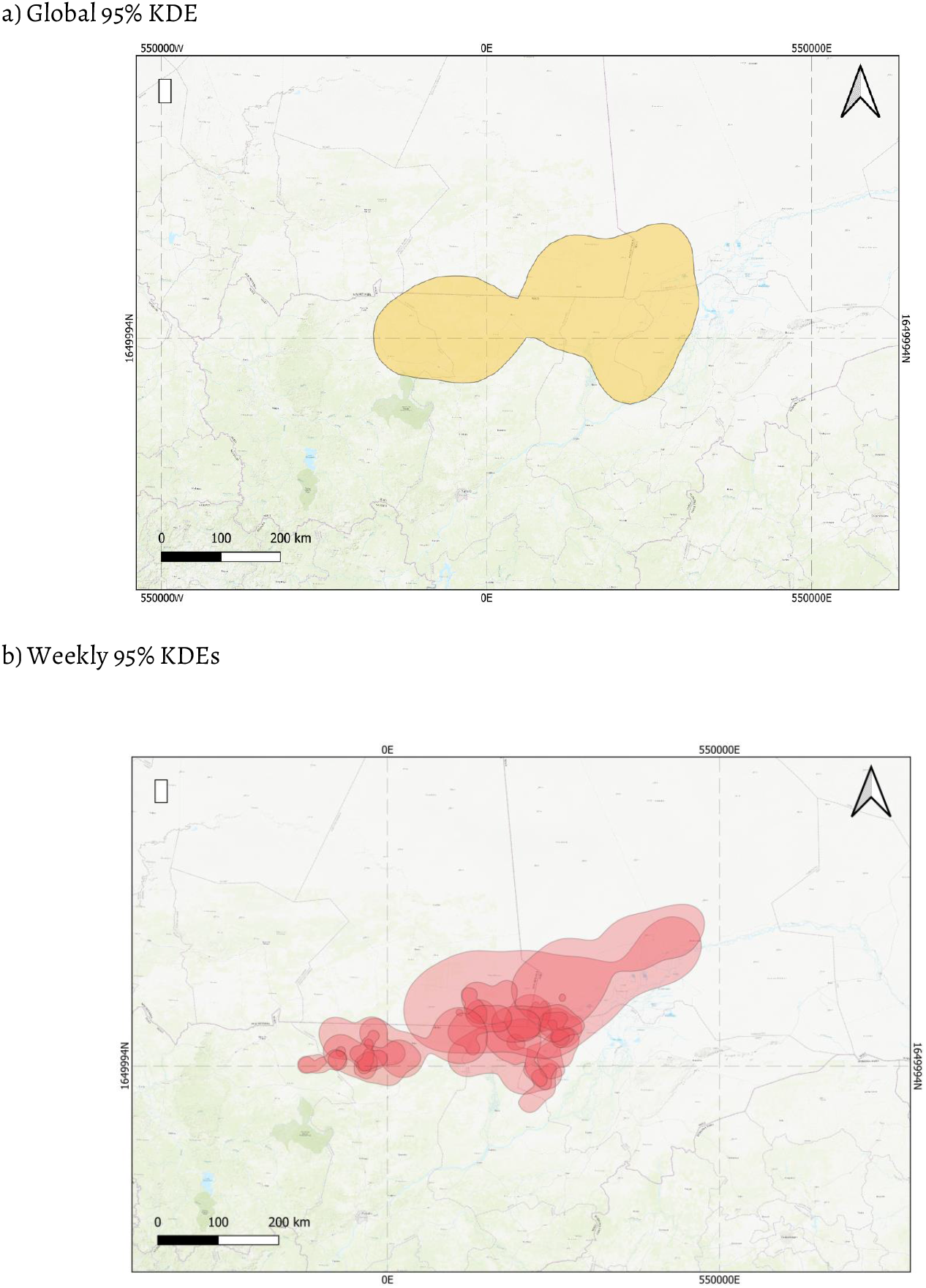
Example of the area occupied by a lesser kestrel (16643) during a single non-breeding period. a) Global 95% KDE, considering all non-breeding locations; b) Weekly 95% KDEs. Each poligon corresponds to a weekly 95% KDE, calculated with the location of each week

**Figure 3.**
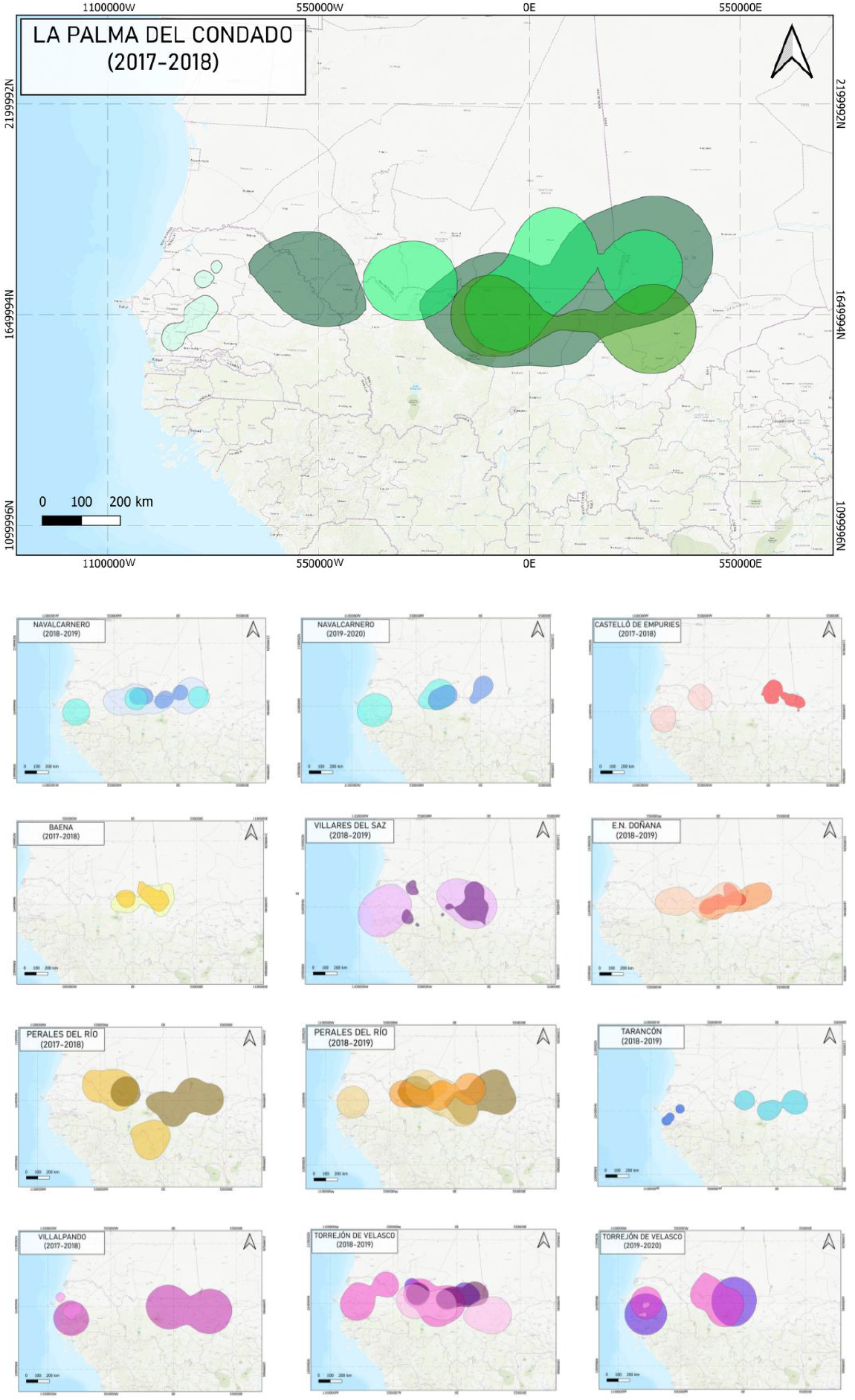
Overlap of non-breeding areas of individuals from the same breeding colony. Each panel represents the overlapping of the members of the colony during a single non-breeding period. Each individual non-breeding area is represented in a different colour.

**Figure 4.**
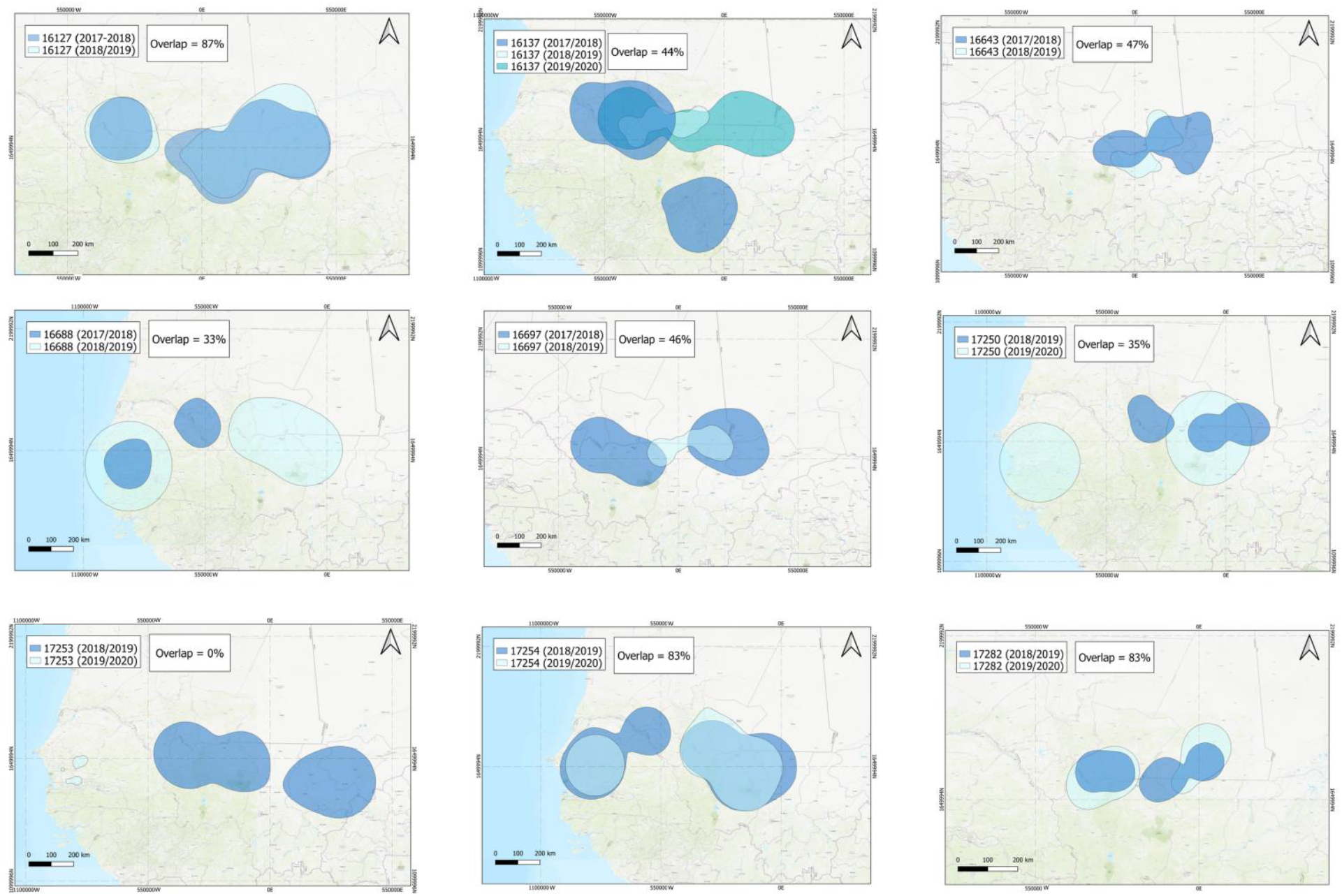
Year-to-year overlap between non-breeding areas of nine individuals. Average overlapping is shown in each panel. Each colour shows a different year.

Individuals traveled, on average, 6205 ± 2407 km during the entire non-breeding period, and 41.1 ± 11.8 km daily (Table 2).

We did not find significant differences between sexes in the size of the non-breeding area or total traveled distances (Table S2), but females tend to cover more daily distances than males (44 vs 38.6 km per day; Table 2).

### Degree of overlap of kestrels of the same colony

Considering each year separately, the non-breeding areas of the individuals from the same breeding colony overlapped by 38.8 ± 21.4 % (range = 0-69). On the other hand, those same areas overlapped with the areas of individuals from other colonies by 40.3 ± 9.5 % (range = 28-61; Table 3). The chi-square test found no significant difference between them (P = 0.9).

**Table 3.**
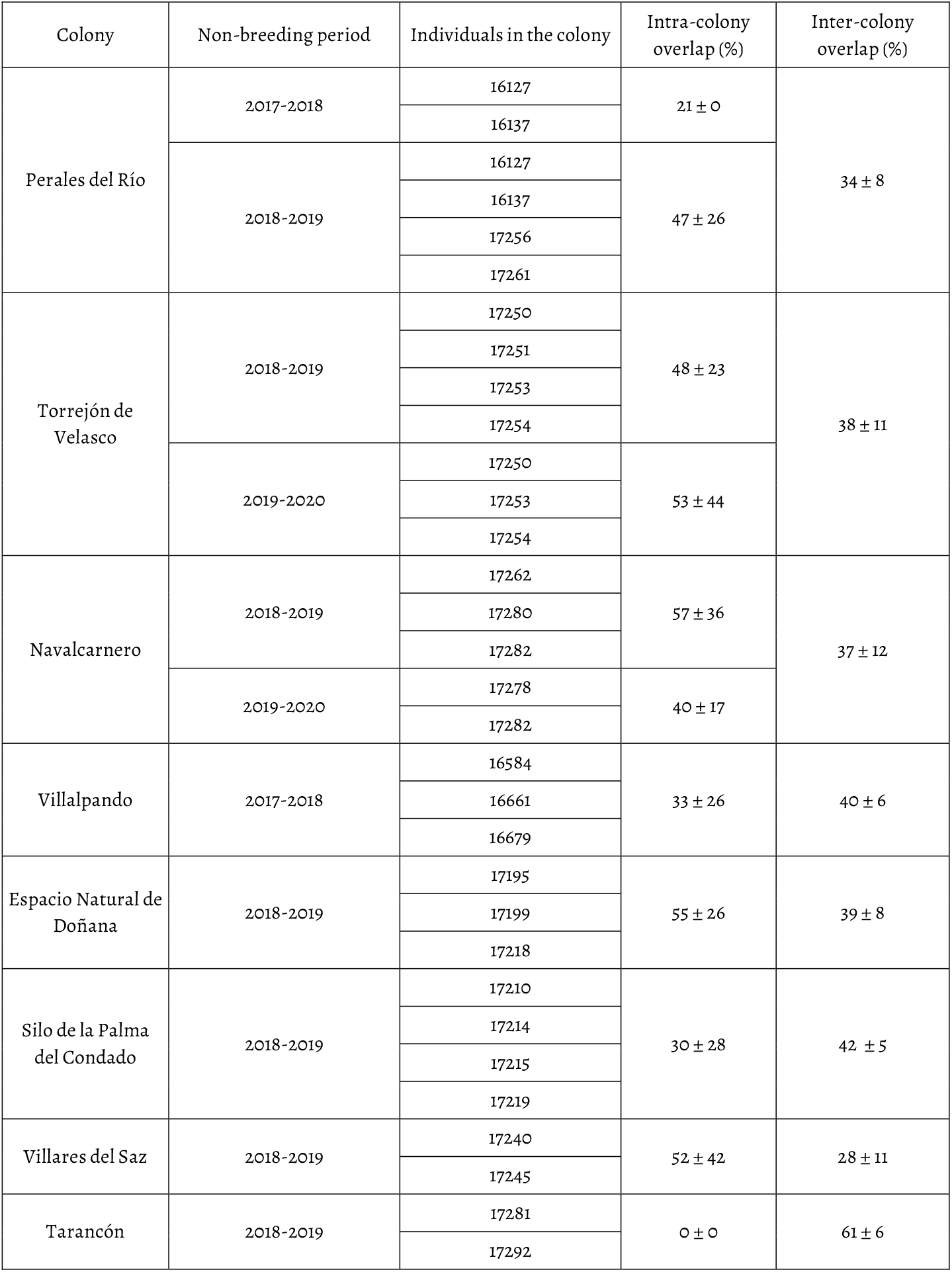
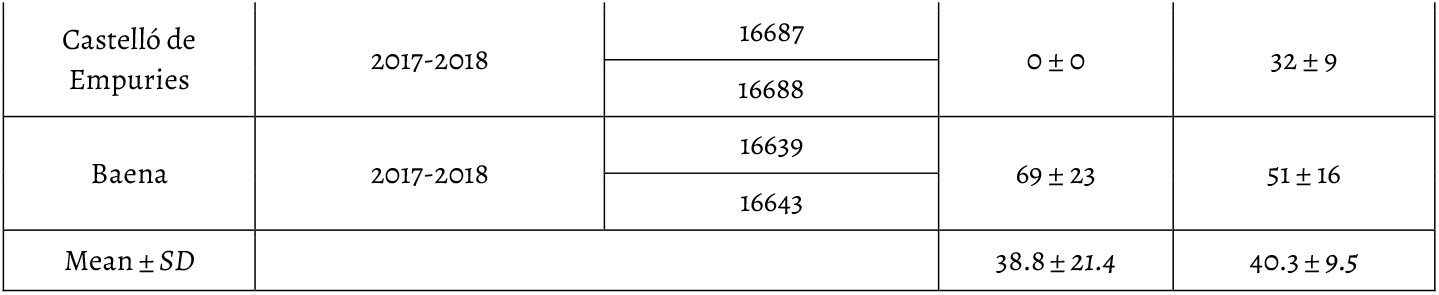
Percentage of overlap between the non-breeding areas (95% KDE) of individuals from the same breeding colony during the same year (intra-colony overlap) and individuals from each breeding colony with individuals from other colonies (inter-colony overlap).

Moreover, all but one individual (8 out of 9) overlapped their non-breeding areas (95% KDE) in consecutive years. The year-to-year overlap of those non-breeding areas was 49 ± 27 % (range = 33-87). Only one individual (17253) changed its non-breeding area, resulting in zero overlap (Table 4).

**Table 4.** Year-to-year overlap of the non-breeding areas (95% KDE) of the same individual (n=9) in consecutive years.

| ID | Year-to-year overlap (%) between non-breeding areas<br>(mean ± SD) |
| --- | --- |
| 16127 | 87 ± 2 |
| 16137 | 44 ± 30 |
| 16643 | 47 ± 26 |
| 16688 | 33 ± 27 |
| 16697 | 46 ± 40 |
| 17250 | 35 ± 22 |
| 17253 | 0 ± 0 |
| 17254 | 83 ± 17 |
| 17282 | 65 ± 10 |
| AVERAGE | 49 ± 27 |

## 4. Discussion

In this work, it is pointed out that lesser kestrels of the same breeding colony do not winter together, but their non-breeding grounds are dispersed from each other. Furthermore, the size of the non-breeding areas, the total and daily traveled distances during this period, and the year-to-year overlap between non-breeding areas of the same individual are studied.

The lesser kestrels used large areas during the non-breeding season (95% KDE = 143,697 ± 98,048 km^2^; range = 5,001-445,393), with no differences between the sexes. Previous work on raptors that spend the non-breeding period in the Sahel also showed non-breeding area sizes within that range: up to 112,000 km^2^ (95% KDE) in the lesser spotted eagles (*Aquila pomarina;* Meyburg *et al*. 2015), 37,508 km^2^ in the Montagu’s harrier (*Circus pygargus;* Limiñana *et al*. 2012b), 26,615 km^2^ in the Egyptian vulture (*Neophron percnopterus;* García-Ripollés *et al*. 2010), or 6,218 km^2^ in the booted eagle (*Aquila pennata*; Urios *et al*. 2017). Limiñana *et al*. (2012a) estimated an area of up to 43,856 km^2^ for the lesser kestrel, which is within the estimated range for the individuals in our study. The large sizes of the non-breeding areas we estimated may respond to different reasons. First, it may be due to the characteristic non-breeding movements of the lesser kestrel. While some species that overwinter in the Sahel are located in a single area which they use throughout the entire season, the lesser kestrels have an itinerant behavior, that is, they move along the longitudinal Sahelian strip and occupy several smaller areas (López-Ricaurte *et al*. 2022). In fact, our weekly estimations revealed areas much smaller (from 64 to 7,359 km^2^) which individuals used to be displaced lengthwise, which also supports the hypothesis of the itinerant behavior during non-breeding season. This behavior could be motivated by the insectivorous diet of the species (Rodríguez *et al*. 2006, Martín *et al*. 2007, Rodríguez *et al*. 2010), which requires greater mobility to provide individuals with sufficient food in winter. On the other hand, the estimations using all the locations of the non-breeding period may overestimate the size of the non-breeding area in the case of itinerant behaviours. Limiñana *et al*. (2012b) discarded the locations between those single areas, which may have caused the area estimation to be reduced. The method to estimate the size of the non-breeding area size should be considered for more detailed comparisons.

The lesser kestrel is a strict colonial species during breeding (Cecere *et al*. 2018), but our results showed that individuals from the same breeding colony overwinter in different areas in the Sahel. Indeed, the overlap between the non-breeding areas of individuals from the same breeding colony is not higher than that expected at random. Previous studies have demonstrated the migratory connectivity of different colonies in Europe: Iberian birds migrate to the western Sahel, Italian birds to the central Sahel, and Balkan birds to the central-eastern Sahel (Sarà *et al*. 2019). Therefore, following a big-scale approach, there is a high degree of connectivity in the non-breeding destinations. However, the more finer-scale approach we tested here showed that individuals from the same breeding colony are not associated during wintering. This phenomenon occurs in other insectivorous birds that winter in the Sahel. For example, for the Montagu’s harriers it has been shown that individuals from the same breeding colony disperse into different non-breeding areas in Africa, up to 1,200 km away. This behaviour was expected because the ecological and reproductive requirements of individuals during the non-breeding period are very different from those during the breeding season (Urios and García-Macía 2022). Without being tied to a nest and a colony, lesser kestrels can optimize territory exploration and foraging by separating non-breeding areas from their colonial partners.

In accordance with previous tracking studies on the Lesser Kestrel during the non breeding period, the repeatability of non-breeding areas was high in the individuals in our study (López-Ricaurte et al., 2022). Lesser kestrels tend to overwinter in the same region than in previous years, which is supported by our results on the year-to-year overlap of non-breeding areas, which ranged from 33 to 87% (49 ± 27 on average). Wintering site fidelity is common among migratory raptors. The experience in the territory obtained over the years allows better energy efficiency and reduces possible risks during non-breeding period (García-Ripollés *et al*. 2010, García-Macía *et al*. 2022).

In conclusion, our study explored the non-breeding behavior of the colonies of lesser kestrels in the Sahel. Their behaviour during non-breeding period tends to be itinerant, so their non-breeding areas are often large and cover a big part of the western Sahel. Individuals from the same breeding colony did not overlap their non-breeding areas more than expected by chance, so apparently the individuals, which were closely spatially linked during the breeding season, selected independent non-breeding areas. On the other hand, there was a high year-on-year repeatability of the non-breeding areas, which allows a better use of the space because of the knowledge of the territory.

## Acknowledgments

We thank M. Aguilera, E. Aguirre, E. Álvarez, P. Aycart, M. Baena, S. Bondì, F. Carbonell, M.A. Carrero, S. De la Fuente, V. De la Torre, M. Galán, M. Garcés, J.L. González, E. Griffin, L. Hernández, E. Holroyd, D. Jordano, P. Lazo, C. Marfil, J. Marín, F.J. Martín-Barranco, R. Mascara, A. Meijide, P. Moreno, D. Ni Dhubhail, C. Ordóñez, M. Pomarol, F.J.Pulpillo, P. Ruiz, A. Valverde. and L. Zanca. for their help during fieldwork and for technical .We especially thank M. Vázquez for his support during fieldwork in Doñana.

This paper is part of Jorge García-Macía’s PhD thesis.

## Funding

Funding for lesser kestrels tagging was provided by Iberdrola España Foundation (MIGRA program of SEO/BirdLife), GREFA (supported by Ministerio para la Transición Ecológica y Reto Demográfico, Junta de Castilla-La Mancha and SEITT, s.a.), Córdoba Zoo, Alcalá de Henares Municipality, and Global Nature Foundation within the LIFE Project “Steppe Farming” (LIFE15417NAT/ES/000734). Lina Lopez-Ricaurte has received financial support through the “La Caixa” INPhINIT Fellowship Grant for Doctoral studies at Spanish Research Centres of Excellence, “La Caixa” Banking Foundation, Barcelona, Spain. In Andalucia tags were funded by “KESTRELS MOVE” project (ref: CGL2016 79249 P) (AEI/FEDER, UE). At the time of analyses and writing, this study was supported by projects MERCURIO (ref: PID2020-421115793GB) (AEI/FEDER,UE) and SUMHAL (European Regional Development Fund4 (ref: LIFEWATCH-2019-09-CSIC-13) (MICINN, POPE 2014-2020). Logistic and technical support was provided by ICTS-RBD.

## Notes

### Competing Interest Statement

The authors have declared no competing interest.

## REFERENCES

Aparicio JM, Bonal R, Muñoz A, 2007. Experimental test on public information use in the colonial lesser kestrel. Evol Ecol 21: 783–800. https://doi.org/10.1007/s10682-006-9151-7

Arroyo B, García J, 2007. El aguilucho cenizo y el aguilucho pálido en España. Población en 2006 y método de censo. Madrid: SEO/BirdLife.

Arroyo B, Mougeot F, Bretagnolle V, 2001. Colonial breeding and nest defence in Montagu’s harrier (*Circus pygargus*). Behav Ecol Sociobiol 50(2): 109–115. https://www.jstor.org/stable/4601942#metadata_info_tab_contents

Barta Z, Giraldeau LA, 2001. Breeding colonies as information centers: a reappraisal of information-based hypotheses using the producer—scrounger game. Behav Ecol 12: 121–127. https://doi.org/10.1093/beheco/12.2.121

Bijleveld M, 1974. Birds of prey in Europe. United Kingdom: MacMillan Press.

Birkhead TR, Møller AP, 1992. Sperm competition in birds. Evolutionary causes and consequences. London: Academic Press.

Bustamante J, Molina B, Del Moral JC, 2020. El cernícalo primilla en España, población reproductora en 2016-18 y método de censo. Madrid: SEO/BirdLife. https://seo.org/boletin/seguimiento/censos/53%20primilla/pdf/SEO%2053%20primilla.pdf

Bustamante J, Molina B, Del Moral JC, 2021. Cernícalo primilla: Falco naumanni. In: López-Jiménez N., editor. Libro Rojo de las Aves de España. Madrid: SEO/BirdLife, 125–136. https://seo.org/wp-content/uploads/2021/12/Libro-Rojo-de-las-Aves-de-Espana-2021.pdf

Cade TJ, Digby RD, 1982. The falcons of the world. USA: Harper Collins.

Cecere JG, Bondì S, Podofillini S, Imperio S, Griggio M, Fulco E, Curcio A, Ménard D, Mellone U, Saino N, Serra L, Sarà M, Rubolini D, 2018. Spatial segregation of home ranges between neighbouring colonies in a diurnal raptor. Sci Rep 8: 11762. https://doi.org/10.1038/s41598-018-29933-2

Clode D, Birks JD, Macdonald DW, 2000. The influence of risk and vulnerability on predator mobbing by terns (*Sterna* spp.) and gulls (*Larus* spp.). J Zool 252: 53–59. https://doi.org/10.1111/j.1469-7998.2000.tb00819.x

Cramp S, Simmons KEL, 1980. Handbook of the Birds of Europe the Middle East and North Africa. The Birds of the Western Palearctic. Vol. II. Hawks to Bustards. United Kingdom: Oxford University Press.

Danchin E, Giraldeau LA, Valone TJ, Wagner RH, 2004. Public information: from nosy neighbors to cultural evolution. Science 305: 487–491. https://doi.org/10.1126/science.1098254

Di Maggio R, Campobello D, Sarà M, 2013. Nest aggregation and reproductive synchrony promote lesser kestrel *Falco naumanni* seasonal fitness. J Ornithol 154: 901–910. https://doi.org/10.1007/s10336-013-0954-3

Ferguson-Lees J, Christie DA 2001. Raptors of the World. United Kingdom: Christopher Helm.

García-Macía J, De La Puente J, Bermejo-Bermejo A, Raab R, Urios V, 2022. High Variability and Dual Strategy in the Wintering Red Kites (*Milvus milvus*). Diversity 14: 117. https://doi.org/10.3390/d14020117

García-Ripollés C, López-López P, Urios V, 2010. First description of migration and wintering of adult Egyptian Vultures *Neophron percnopterus* tracked by GPS satellite telemetry. Bird Study 57(2): 261–265, https://doi.org/10.1080/00063650903505762

GBIF, 2022. Falco naumanni Fleischer, 1818. https://www.gbif.org/es/species/9584698

Grainger Hunt W, Dunlop N, Lockhart JM, 2021. A revealing case of territorial fighting by Golden Eagles. J Raptor Res 55(1): 112–114. https://doi.org/10.3356/0892-1016-55.1.112

IUCN, 2021. The IUCN Red List of Threatened Species. Version 2021-3. Available at: www.iucnredlist.org. (Accessed: 09 December 2021).

Kenward RE, 2001. A manual for wildlife radio tagging. London: Academic Press.

Lack D, 1968. Ecological adaptations for breeding in birds. London: Chapman & Hall.

Limiñana R, Romero M, Mellone U, Urios V, 2012a. Mapping the migratory routes and wintering areas of lesser kestrels *Falco naumanni:* new insights from satellite telemetry. Ibis 154(2): 389–399. https://doi.org/10.1111/j.1474-919X.2011.01210.x

Limiñana R, Soutullo A, Urios V, Reig-Ferrer A, 2012b. Migration and wintering areas of adult Montagu’s Harriers (*Circus pygargus*) breeding in Spain. J Ornithol 153: 85–93. https://doi.org/10.1007/s10336-011-0698-x

López-López P, Perona A, Egea-Casas O, Etxebarria Morant J, Urios V, 2021. Tri-axial accelerometry shows differences in energy expenditure and parental effort throughout the breeding season in long-lived raptors. Curr Zool 1: 57–67. https://doi.org/10.1093/cz/zoab010

López-Ricaurte L, Wouter MGV, Hernández-Pliego J, García-Silveira D, Casado S, Garcés-Toledano F, Martínez-Dalmau J, Ortega A, Rodríguez-Moreno B & Bustamante J. (In review). Itinerant lifestyle and congregation of lesser kestrels in West Africa. https://doi.org/10.1101/2022.08.12.503182

Martín M, Guerrero M, Mendoza P, Antolín J, 2007. Experiencia con cámara web para la determinación del régimen alimenticio en la ZEPA “Iglesia de la Purificación” de Almendralejo extremadura. Primilla info 6: 11–13.

Meyburg BU, Mendelsohn S, Mendelsohn J, De Klerk HM, 2015. Revealing unexpected uses of space by wintering *Aquila pomarina:* How does satellite telemetry identify behaviour at different scales? J Avian Biol 46: 648–657. https://doi.org/10.1111/jav.00670

Ortego J, 2016. Cernícalo primilla – Falco naumanni. In: Salvador A, Morales MB, editors. Enciclopedia Virtual de los Vertebrados Españoles. Madrid: Museo Nacional de Ciencias Naturales. http://www.vertebradosibericos.org/

Pérez-García JM, Margalida A, Afonso I, Ferreiro E, Gardiazábal A, Botella F, Sánchez-Zapata JA, 2013. Interannual home range variation, territoriality and overlap in breeding Bonelli’s Eagles (*Aquila fasciata*) tracked by GPS satellite telemetry. J Ornithol 154: 63–71. https://doi.org/10.1007/s10336-012-0871-x

Pilard P, Lelong V, Sonko A, Riols C, 2011. Suivi et conservation du dortoir de rapaces insectivores (*Faucon crécerellette Falco naumanni* et Elanion Naucler *Chelictinia riocouriĩ*) de l’Ile de Kousmar (Kaolack/Sénégal). Alauda 79: 295–312.

Rodríguez C, Johst K, Bustamante J, 2006. How do crop types influence breeding success in lesser kestrels through prey quality and availability? A modelling approach. J Appl Ecol 43(3): 587–597. https://www.jstor.org/stable/3838461

Rolland C, Danchin E, de Fraipont M, 1998. The evolution of coloniality in birds in relation to food, habitat, predation, and life-history traits: A comparative analysis. Am Nat, 151(6): 514–529. https://doi.org/10.1086/286137

Rodríguez C, Tapia L, Kieny F, Bustamante J, 2010. Temporal changes in lesser kestrel (*Falco naumanni*) diet during the breeding season in southern Spain. J Raptor Res 44(2): 120–128. https://doi.org/10.3356/JRR-09-34.1

Sarà M, Bondi S, Bermejo A, Bourgeois M, Bouzin M, Bustamante J,… Rubolini D, 2019. Broad-front migration leads to strong migratory connectivity in the lesser kestrel (*Falco naumanni*). J Biogeograpr 46(12): 2663–2677. https://doi.org/10.1111/jbi.13713

Senar JC, Borras A, 2004. Sobrevivir al invierno: estrategias de las aves invernantes en la Península Ibérica. Ardeola, 51 (1): 133–168. https://www.ardeola.org/uploads/articles/docs/557.pdf

Urios V, Bermejo A, Vidal-Mateo J, De la Puente J, 2017. Migración y ecología espacial de la población española de águila calzada. Madrid: SEO/BirdLife.

Urios V, García-Macía J, 2022. Migración y ecología espacial de la población española de milano real. Madrid: SEO/BirdLife. https://seo.org/boletin/seguimiento/migracion/08_milano_real/08_milano_real.pdf

Ward P, Zahavi A, 1973. The importance of certain assemblages of birds as “information-centres” for food-finding. Ibis 115: 517–534. https://doi.org/10.1111/j.1474-919X.1973.tb01990.x

Wittenberger JF, Hunt GL, 1985. The adaptive significance of coloniality in birds. Avian Biol, 8: 1–78. https://doi.org/10.1016/B978-0-12-249408-6.50010-8

