## Supplementary Materials for "Kestrels of the same colony do not overwinter together"

Table S1. Values of the 55 non-breeding periods, regarding dates, duration, travelled distances and kernel density estimations (KDE). Daily travelled distances are shown as "mean ± *SD* (minimum-maximum)". KDE = Kernel Density Estimation.

| ID | Sex | Non-breeding period | Beggining of non-breeding period | End of non-breeding period | Days of non-breeding period | Total travelled distance (km) | Daily travelled distance (km) | Global 95% KDE (km²) | Global 50% KDE (km²) | Median 95% KDE (km²) | Median 50% KDE (km²) |
|---|---|---|---|---|---|---|---|---|---|---|---|
| 16127 | Male | 2017/2018 | 42986 | 43157 | 171 | 8706 | 50.9 ± 39.9 (0.4-235.1) | 228443 | 49861 | 3073 | 810 |
| | | 2018/2019 | 43370 | 43525 | 155 | 6231 | 40.3 ± 35.2 (0.1-171.0) | 234612 | 47788 | 1861 | 344 |
| 16137 | Male | 2017/2018 | 42998 | 43193 | 195 | 11039 | 56.7 ± 48.3 (0.3-268.3) | 224158 | 49205 | 2570 | 553 |
| | | 2018/2019 | 43390 | 43527 | 137 | 4178 | 30.6 ± 31.8 (0.04-149.3) | 44429 | 8326 | 7359 | 1403 |
| | | 2019/2020 | 43739 | 43882 | 143 | 6245 | 43.7 ± 40.2 (0.1-247.1) | 192739 | 41289 | 470 | 68 |
| 16212 | Male | 2016/2017 | 42642 | 42798 | 155 | 10030 | 64.5 ± 48.4 (0.02-127.9) | 248652 | 56552 | 1591 | 335 |
| 16275 | Female | 2017/2018 | 43006 | 43183 | 177 | 10401 | 58.9 ± 46.3 (0.1-215.7) | 227373 | 47972 | 1095 | 164 |
| 16584 | Female | 2017/2018 | 42985 | 43187 | 202 | 7642 | 37.9 ± 44.4 (0.1-286.4) | 312294 | 58035 | 2115 | 375 |
| 16611 | Male | 2017/2018 | 42988 | 43183 | 195 | 10000 | 51.3 ± 45.1 (0.1-252.2) | 49480 | 11686 | 5066 | 1175 |
| | | 2018/2019 | 43374 | 43536 | 162 | 6228 | 38.5 ± 39.3 | 67728 | 16967 | 4355 | 658 |

| | | | | | | | | | | | |
|---|---|---|---|---|---|---|---|---|---|---|---|
| | | | | | | | (0.1-171.8) | | | | |
| 16616 | Female | 2017/2018 | 42982 | 43029 | 47 | 1112 | 23.6 ± 51.8 (0.2-271.4) | 125283 | 16610 | 1807 | 268 |
| 16639 | Male | 2017/2018 | 42999 | 43149 | 150 | 6927 | 46.3 ± 37.2 (0.8-204.7) | 58402 | 11000 | 2435 | 453 |
| 16643 | Female | 2017/2018 | 43004 | 43154 | 150 | 9024 | 60.3 ± 45.0 (0.5-211.2) | 95317 | 24046 | 2437 | 464 |
| | | 2018/2019 | 43395 | 43512 | 117 | 4036 | 34.6 ± 33.1 (1.3-183.9) | 42793 | 9169 | 1012 | 186 |
| 16661 | Female | 2017/2018 | 42992 | 43165 | 173 | 8650 | 49.9 ± 41.4 (0.03-165.2) | 21983 | 4631 | 793 | 180 |
| 16679 | Female | 2017/2018 | 42987 | 43172 | 185 | 10600 | 57.4 ± 43.7 (0.8-209.1) | 180889 | 38867 | 835 | 118 |
| | | 2018/2019 | 43383 | 43534 | 151 | 9548 | 63.3 ± 46.2 (0.6-299.4) | 42277 | 8536 | 1174 | 196 |
| 16687 | Female | 2017/2018 | 43016 | 43172 | 156 | 9336 | 60.0 ± 54.6 (2.5-342.9) | 44684 | 7675 | 1336 | 266 |
| 16688 | Male | 2017/2018 | 43016 | 43166 | 150 | 6595 | 44.0 ± 32.2 (0.6-258.4) | 72539 | 15192 | 350 | 59 |
| | | 2018/2019 | 43394 | 43527 | 133 | 3547 | 26.7 ± 29.6 (0.5-183.9) | 274837 | 58163 | 2233 | 424 |
| 16690 | Male | 2017/2018 | 43013 | 43209 | 196 | 7648 | 39.0 ± 40.4 (0.6-201.7) | 74595 | 8589 | 253 | 37 |
| | | 2018/2019 | 43394 | 43526 | 132 | 4301 | 32.6 ± 48.0 (0.01-261.22) | 87623 | 17210 | 608 | 102 |
| 17195 | Male | 2018/2019 | 43387 | 43463 | 76 | 1840 | 24.4 ± 25.7 (0.4-116.9) | 152104 | 32533 | 1702 | 310 |
| 17199 | Female | 2018/2019 | 43383 | 43506 | 123 | 4027 | 32.8 ± 30.7 (1.8-153.4) | 50030 | 10367 | 333 | 75 |

| | | | | | | | | | | | |
|---|---|---|---|---|---|---|---|---|---|---|---|
| 17210 | Male | 2018/2019 | 43400 | 43498 | 98 | 3901 | 39.8 ± 42.1 (2.3-281.8) | 92847 | 16668 | 982 | 163 |
| 17214 | Female | 2018/2019 | 43365 | 43506 | 141 | 5381 | 38.2 ± 35.8 (2.4-197.2) | 155202 | 38026 | 3105 | 434 |
| 17215 | Female | 2018/2019 | 43377 | 43525 | 148 | 7410 | 50.0 ± 43.0 (1.9-263.3) | 263758 | 44396 | 1218 | 165 |
| 17218 | Male | 2018/2019 | 43390 | 43512 | 122 | 3921 | 32.2 ± 43.0 (0.9-201.7) | 105025 | 24343 | 1294 | 236 |
| 17219 | Male | 2018/2019 | 43380 | 43530 | 150 | 6042 | 40.3 ± 29.2 (1.8-143.7) | 12412 | 1782 | 409 | 62 |
| 17235 | Male | 2018/2019 | 43364 | 43525 | 161 | 5568 | 34.7 ± 41.4 (0.1-272.5) | 51212 | 9622 | 657 | 111 |
| | | 2019/2020 | 43731 | 43863 | 132 | 3860 | 29.3 ± 34.4 (0.5-170.8) | 232516 | 40142 | 5021 | 855 |
| 17237 | Male | 2018/2019 | 43392 | 43526 | 134 | 6301 | 47.0 ± 34.7 (3.2-253.7) | 111628 | 25882 | 417 | 60 |
| | | 2019/2020 | 43738 | 43887 | 149 | 5627 | 37.9 ± 30.2 (0.2-149.4) | 279949 | 50695 | 942 | 172 |
| 17239 | Female | 2018/2019 | 43374 | 43517 | 143 | 8592 | 60.2 ± 51.2 (0.5-329.6) | 280088 | 53519 | 245 | 28 |
| | | 2019/2020 | 43737 | 43887 | 150 | 4188 | 28.0 ± 28.8 (0.04-214.6) | 100827 | 21004 | 1331 | 190 |
| 17240 | Male | 2018/2019 | 43387 | 43539 | 152 | 4930 | 32.4 ± 43.0 (1.3-253.6) | 86724 | 11163 | 659 | 132 |
| 17241 | Female | 2018/2019 | 43731 | 43905 | 173 | 7862 | 45.3 ± 41.6 (0.1-195.9) | 120676 | 18843 | 517 | 85 |
| 17245 | Female | 2018/2019 | 43383 | 43517 | 134 | 6909 | 51.5 ± 53.7 (1.2-308.9) | 309407 | 67822 | 1534 | 238 |
| 17250 | Male | 2018/2019 | 43370 | 43513 | 143 | 3981 | 27.9 ± 36.4 | 90574 | 20060 | 112 | 19 |

| | | | | | | | | | | | |
|---|---|---|---|---|---|---|---|---|---|---|---|
| | | | | | | | (1.0-158.3) | | | | |
| | | 2019/2020 | 43729 | 43879 | 150 | 4625 | 30.9 ± 50.1 (0.2-451.5) | 239308 | 51537 | 711 | 77 |
| 17251 | Male | 2018/2019 | 43373 | 43503 | 130 | 5458 | 42.0 ± 33.1 (2.3-153.5) | 109293 | 27746 | 517 | 108 |
| 17253 | Male | 2018/2019 | 43401 | 43526 | 125 | 3468 | 27.8 ± 31.8 (1.3-231.3) | 227611 | 46382 | 1425 | 265 |
| | | 2019/2020 | 43749 | 43886 | 137 | 2923 | 21.4 ± 21.3 (0.1-106.2) | 5001 | 750 | 76 | 15 |
| 17254 | Male | 2018/2019 | 43369 | 43526 | 157 | 6198 | 39.6 ± 42.5 (0.1-271.4) | 269755 | 52646 | 358 | 78 |
| | | 2019/2020 | 43730 | 43872 | 142 | 8034 | 56.4 ± 52.4 (2.4-325.0) | 202624 | 43147 | 1650 | 297 |
| 17256 | Female | 2018/2019 | 43412 | 43528 | 116 | 4616 | 39.9 ± 37.4 (1.9-176.4) | 158512 | 35479 | 1287 | 227 |
| 17261 | Male | 2018/2019 | 43368 | 43526 | 158 | 4724 | 29.9 ± 44.5 (1.9-343.9) | 230772 | 53674 | 125 | 23 |
| 17262 | Female | 2018/2019 | 43377 | 43526 | 149 | 5296 | 35.6 ± 37.8 (0.4-217.6) | 182647 | 35185 | 231 | 41 |
| 17278 | Female | 2019/2020 | 43725 | 43893 | 167 | 9506 | 56.8 ± 45.3 (1.1-324.7) | 126928 | 25837 | 487 | 71 |
| 17280 | Female | 2018/2019 | 43370 | 43526 | 156 | 7127 | 45.8 ± 49.2 (1.2-230.3) | 445393 | 96323 | 317 | 73 |
| 17281 | Female | 2018/2019 | 43374 | 43526 | 152 | 6511 | 42.9 ± 36.7 (0.9-188.9) | 86870 | 18184 | 64 | 7 |
| 17282 | Female | 2018/2019 | 43379 | 43511 | 132 | 2649 | 20.1 ± 26.5 (0.9-145.3) | 54889 | 12742 | 210 | 39 |
| | | 2019/2020 | 43724 | 43861 | 137 | 3315 | 24.2 ± 26.3 (2.5-136.2) | 68249 | 14296 | 1135 | 167 |

| | | | | | | | | | | | |
|---|---|---|---|---|---|---|---|---|---|---|---|
| 17288 | Female | 2018/2019 | 43363 | 43537 | 174 | 5676 | 32.7 ± 31.9 (0.1-167.7) | 36586 | 6842 | 739 | 145 |
| 17292 | Female | 2018/2019 | 43370 | 43541 | 171 | 8772 | 51.4 ± 41.2 (0.2-161.0) | 12770 | 2764 | 92 | 19 |

**Table S2.** Linear Mixed Models (a) and results of Mann-Whitney-Wilcoxon tests (b) on the comparison between sexes.

**a)**

| Model | n | | Estimate | Standard deviation | Degrees of freedom | P-value |
|---|---|---|---|---|---|---|
| **Total travelled distance** ~ Sex + (1\|Individual) + (1\|Year) | 55 | **Intercept** | 7430.17 | 1114.27 | 6.67 | **0.009** |
| | | **Sex (Male)** | -945.46 | 624.55 | 34.09 | 0.139 |
| **Daily travelled distance** ~ Sex + (1\|Individual) + (1\|Year) | 55 | **Intercept** | 47.44 | 5.23 | 1.75 | **0.0179** |
| | | **Sex (Male)** | -5.19 | 3.06 | 36.32 | 0.098 |

**b)**

| Comparison | n | W | P-valor |
|---|---|---|---|
| **Total 95% KDE ~ Sex** | 55 | 348 | 0.657 |
| **Total 50% KDE ~ Sex** | 55 | 335 | 0.507 |
| **Median weekly 95% KDE ~ Sex** | 55 | 335 | 0.507 |
| **Median weekly 50% KDE ~ Sex** | 55 | 334 | 0.497 |
